# Atomic-level architecture of *Caulobacter crescentus* flagellar filaments provide evidence for multi-flagellin filament stabilization

**DOI:** 10.1101/2023.07.10.548443

**Authors:** Juan C. Sanchez, Eric J. Montemayor, Nicoleta T. Ploscariu, Daniel Parrell, Joseph K. Baumgardt, Jie E. Yang, Bryan Sibert, Kai Cai, Elizabeth R. Wright

## Abstract

Flagella are dynamic, ion-powered machines with assembly pathways that are optimized for efficient flagella production. In bacteria, dozens of genes are coordinated at specific times in the cell lifecycle to generate each component of the flagellum. This is the case for *Caulobacter crescentus*, but little is known about why this species encodes six different flagellin genes. Furthermore, little is known about the benefits multi-flagellin species possess over single flagellin species, if any, or what molecular properties allow for multi-flagellin filaments to assemble. Here we present an in-depth analysis of several single flagellin filaments from *C. crescentus,* including an extremely well-resolved structure of a bacterial flagellar filament. We highlight key molecular interactions that differ between each bacterial strain and speculate how these interactions may alleviate or impose helical strain on the overall architecture of the filament. We detail conserved residues within the flagellin subunit that allow for the synthesis of multi-flagellin filaments. We further comment on how these molecular differences impact bacterial motility and highlight how no single flagellin filament achieves wild-type levels of motility, suggesting *C. crescentus* has evolved to produce a filament optimized for motility comprised of six flagellins. Finally, we highlight an ordered arrangement of glycosylation sites on the surface of the filaments and speculate how these sites may protect the β-hairpin located on the surface exposed domain of the flagellin subunit.

## Introduction

Bacterial motility is characterized as the ability of a cell to tumble, swim, and swarm from unfavorable conditions towards more favorable conditions, typically those higher in nutrients. Bacteria use a combination of pili and flagella for such coordinated movements. The bacterial flagellum is a dynamic, ion-powered machine that requires the coordinated synthesis of over 60 genes, encompassing both structural and regulatory proteins (1). The flagellum synthesis pathway is fairly conserved throughout bacteria and is tightly regulated due to its high energetic cost (2). In the gram-negative, oligotrophic bacterium *Caulobacter crescentus*, a single polar flagellum is synthesized 30 to 40 minutes before cell division ensuring motility in the resulting new swarmer cell (3).

The flagellum is comprised of a basal body consisting of the MS-ring, C-ring, and rod. The MS-ring is a transmembrane complex that acts as a foundation for the flagellum motor and filament structure (4). The C-ring assembles on the cytoplasmic side of the MS-ring and anchors the associated Type 3 secretion apparatus involved in torque generation and rotation switching (1, 4). Establishment of the MS-ring and C-ring is critical for synthesis of the non-cytoplasmic structures of the flagella, as structural proteins are secreted through a central pore of the MS-ring and are incorporated at the distal end of the nascent flagellum (5). The first non-cytoplasmic structure is the rod, and it serves two purposes in *C. crescentus*, first it acts as a checkpoint that coordinates flagellar gene expression with assembly, and second it breaches the cell surface to begin synthesis of the extracellular structures (1).

Completion of the basal structure is followed by secretion of hook structural proteins that assemble the hook on to the distal end of the rod (6). The hook is a curved filamentous structure spanning ∼55 nm and serves as a universal joint, transferring torque from the motor to the filament (7, 8). Upon completion, hook-associated proteins localize to the distal end of the hook to form a hook-filament junction that serves as a platform for the filament cap protein (9). The filament cap protein is instrumental in the polymerization of tens of thousands of flagellins, the filament structural protein, to the growing flagellar filament and remains at the distal end of the filament during filament elongation (10). Flagellar filaments have been reported to be as long as 15 μm in *Escherichia coli* and *Salmonella* species, while in *C. crescentus* filaments are on average 6 μm in length (10–14).

Some species, such as *E. coli,* form a flagellar filament comprised of a single flagellin type and the paradigm of a symmetric homopolymeric filament was widely accepted. However, 45% of bacterial species encode more than one flagellin, ranging from two (*Salmonella enterica* serovar Typhimurium) to fifteen (*Magnetococcus* sp. MC-1) (15, 16). Additionally, the incorporation of these subunits into the filament varies. For example, *Salmonella enterica* serovar *typhimurium* undergoes phase variation where only one of the two flagellin types is incorporated into the filament (17). In *Campylobacter jejuni* and *Bacillus thuringiensis*, multiple flagellins are encoded and assembled into the filament (18, 19). In *C. crescentus,* six flagellin genes are encoded, synthesized, and incorporated into the nascent filament (15). In addition, wild-type *C. crescentus* filaments exhibit spatiotemporal organization with distinct regions of the filament comprised of certain flagellin types (13, 20). Immuno-electron microscopy studies revealed that a 29 kDa flagellin (FljJ) is located proximal to the hook, followed by a 1-2 μm region of a 27 kDa flagellin (FljL), while the distal end of the filament is comprised of a 25 kDa flagellin (FljK) (20). Later, three additional 25 kDa flagellins, denoted FljM, FljN, and FljO, were identified. These flagellins are encoded on the *C. crescentus* genome and are believed to make up the distal region of the filament, though this has not been confirmed (13, 21). Several combinations of these flagellins form viable flagellated cells, and all single flagellin filaments, studied to date, have reduced motility profiles as compared to the wild-type strain (13, 15). Although the wild-type filament of *C. crescentus* has been optimized for swimming, little is known about the role of varying flagellin subunits in the flagellar filament. In agreement with our observations of multi-flagellin filaments, *Pseudomonas aeruginosa* also exhibits reduction in motility when one of its two flagellin genes are perturbed (22). While in *Bdellovibrio bacteriovorus* the flagellar filament is organized into distinct regions as observed in *C. crescentus* (23). These results suggest that flagellar filaments are much more structurally complex than originally thought.

Cryo-EM has been instrumental in revealing additional details about these macromolecular complexes with the first complete atomic model of the bacterial flagellar filament, from *S. enterica*, detailing a helical assembly comprised of 11 protofilaments with a precise rise and twist per adjacent flagellin monomer (24). The flagellin subunit was determined to consisted of 4 domains, with the inner core domains D0 and D1 comprised of coiled coils and the surface exposed domains D2 and D3 consisting of β-sheets (24). As cryo-EM capabilities improved, further structural insight was gained from structural analysis of *B. subtilis* and *P. aeruginosa* filaments, including how some flagellins lack the outer D2 and D3 domains and how unique interactions between adjacent flagellins govern molecular packing of the filament subunits (25). However, these studies assessed filaments containing point mutations which altered their flagellin packing and locked them into straightened forms (24, 25). As cryo-EM data processing methods improve there is interest in understanding how interactions in non-straightened structures impacts the overall helical architecture of the filament (13, 26, 27).

Here, we present three cryo-EM structures of *C. crescentus* flagellar filaments FljK, FljL, and FljM. Each structure was resolved to a global resolution below 3.0 Å, and we report an extremely well-resolved flagellar filament structure of *C. crescentus* FljM at 2.11 Å. We detail how key molecular interactions along the 11-start, 6-start, and 5-start axis differ between each filament model and impact flagellin packing, altering the overall helical architecture of the filament. Interestingly, we show how specific surface threonine residues are glycosylated and form organized lattices that may protect the surface exposed β-hairpin region. In addition, motility assays reveal that the single flagellin mutants for FljK, FljL and FljM, in this analysis, cannot recover wild-type levels of motility, suggesting the multi-flagellin *C. crescentus* filament is best optimized for swimming.

## Results

### Cryo-EM Helical Reconstructions of FljK, FljL, and FljM Flagellar Filaments

It is well-established that a bacterial flagellum comprised of any number of flagellin sub-units is vital for cell motility. Motility assays of *Caulobacter* strains with mono-flagellin filaments indicated reduced motility when compared to the wild-type strain (Fig. S1) (13). Therefore, using cryo-EM and helical reconstruction methods, we sought to determine how packing and filament organization of individual flagellins produced energetically stable filaments.

Reconstruction of electron-potential maps followed a similar workflow for all three mono-flagellin filament samples (Fig. S2, S4, S6). Key differences with previous *Caulobacter* flagella reconstruction methods included 1) the integration of a Topaz neural network for automated particle picking, 2) expansion of the particle box size to 500 pixels (∼420 nm), 3) and exporting particles from RELION to cryoSPARC for additional 3D classification, asymmetrical reconstruction, and 3D variability analysis (Fig. S2-S7) (28, 29). The resulting maps resolved to below 3 Å and showed clear secondary structures and side-chain densities suitable for model building (Fig. 1, Table 1).

**Figure 1.**
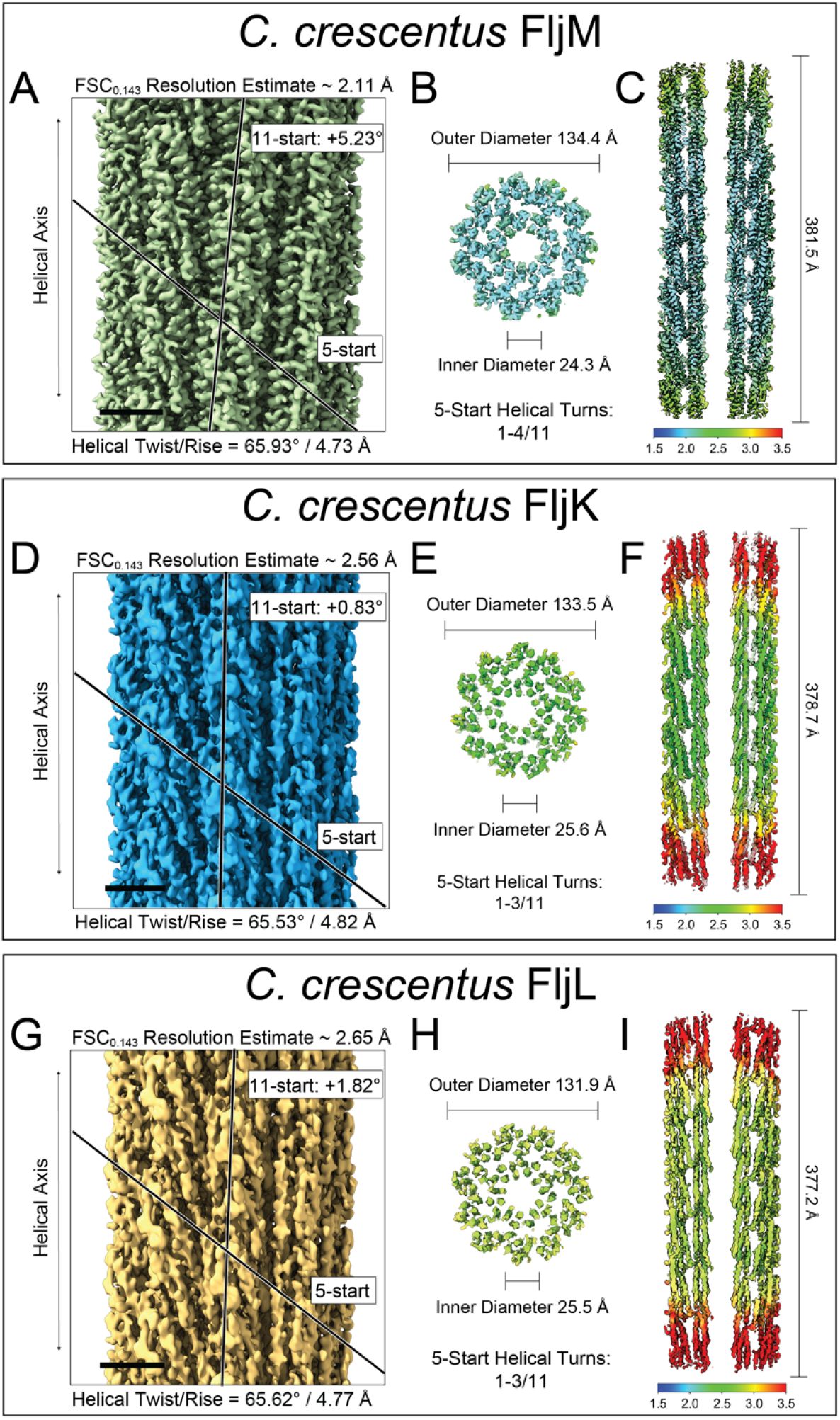
Cryo-EM reconstructions and resolution maps of *C. crescentus* FljM, FljK, and FljL flagellar filaments. (A, D, G) Side view of the *C. crescentus* FljM (A), FljK (D), and FljL (G) helical reconstructions at 2.11 Å, 2.56 Å, and 2.65 Å resolution, respectively (scale bar, 25 Å). (B, E, H) Cross-section of the FljM (B), FljK (E), and FljL (H) filament resolution maps with measurements of the central lumen (inner diameter) and filament (outer diameter). (C, F, I) Cross-section along the helical axis of the FljM (C), FljK (F), and FljL (I) filament resolution map with length measurements of the reconstructions. Resolution heat maps generated in cryoSPARC range from 1.5 Å (blue) to 3.5 Å (red).

**Table 1.**
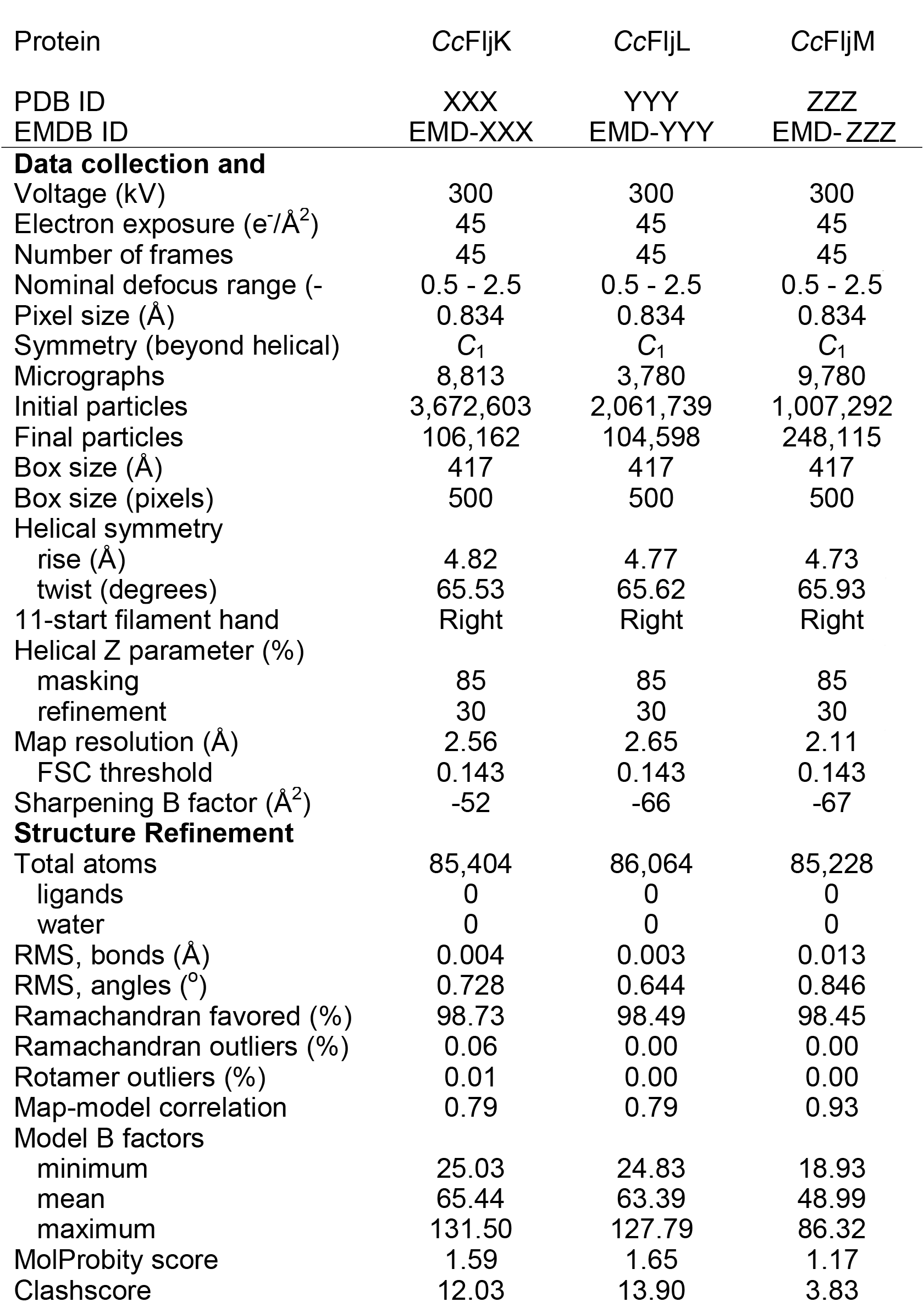
Data collection, symmetrized reconstruction processed in RELION and cryoSPARC, and model statistics

Initial helical parameters for all three reconstructions were obtained from previously solved structures of homologous flagellins (13). These parameters were applied to a featureless cylinder and then optimized in RELION-4.0 and cryoSPARC (Fig. S2-S7). The FljK filament reconstruction resolved to ∼2.56 Å with a helical rise of 4.82 Å and a helical twist of 65.53° (Fig. 1B). The rotational displacement of flagellins along one protofilament, also termed the 11-start angle, was determined to be 0.83° for the FljK filament (Fig. 1B, 2B). Additionally, there was a translational displacement of 53.13 Å along the 11-start between FljK subunits (Fig. 2B). The FljL model resolved to ∼2.65 Å with a helical rise of 4.77 Å and a helical twist of 65.62° (Fig. 1C). The rotational displacement of flagellins along the 11-start angle was 1.83° with a translational displacement of 52.58 Å between intra-protofilament subunits (Fig. 1C, 2C). Finally, the FljM filament was the best resolved flagellum reconstruction to date at ∼2.11 Å, with a helical rise of 4.73 Å and a helical twist of 65.93° (Fig 1 A). The intra-protofilament subunits had a rotational displacement of 5.23° and translational displacement of 52.03 Å (Fig. 2A). Additionally, all three structures displayed the conserved 11 protofilament archetype observed in bacterial flagellum models (Fig. 2).

**Figure 2.**
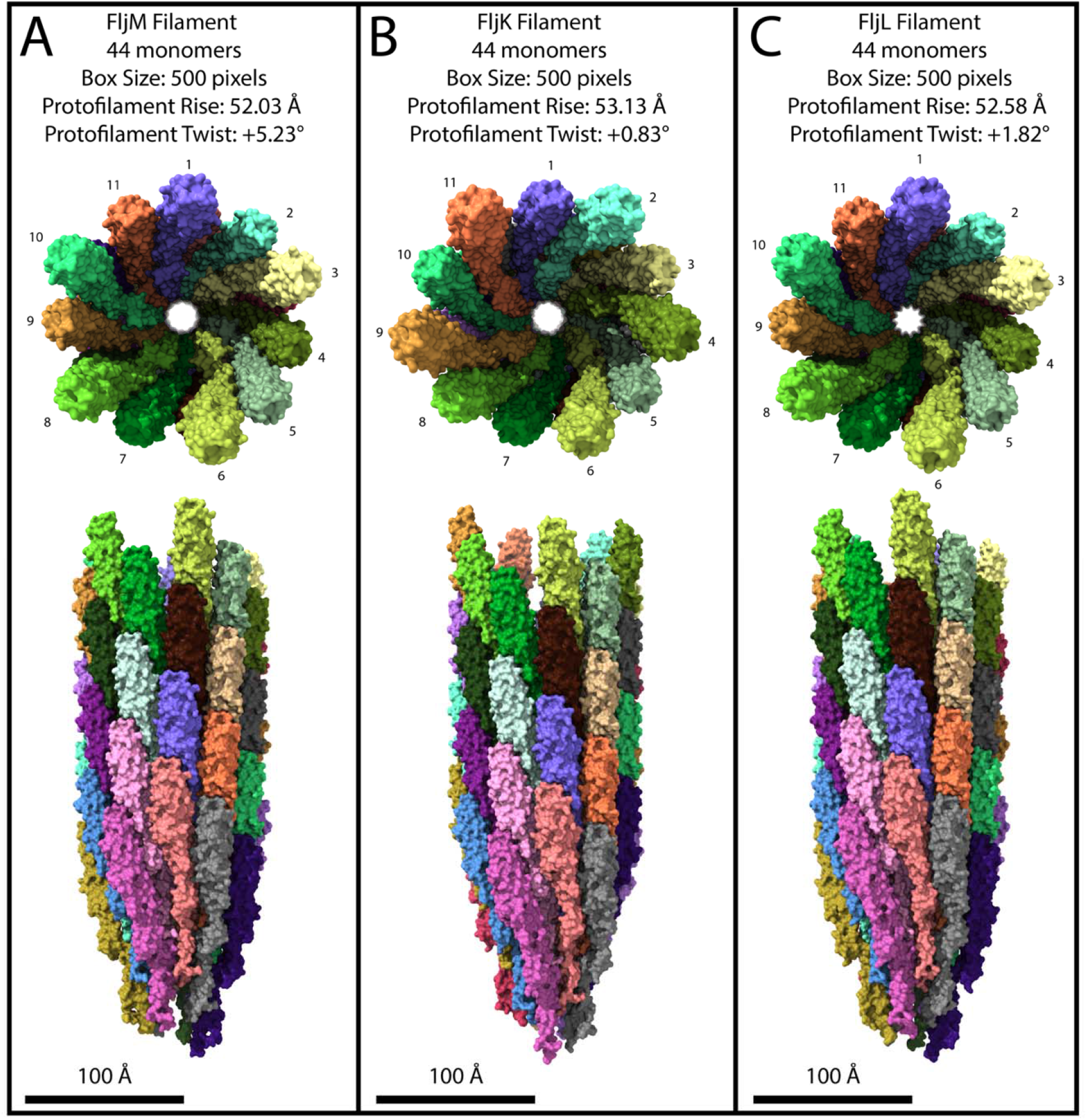
Atomic models of the *C. crescentus* FljM, FljK, and FljL flagellar filaments. Top view and side view of the FljM (A), FljK (B), and FljL (C) filament models built into the corresponding electron potential maps (Figure 1). Top views detail the archetypal 11 protofilament structure (scale bar, 100 Å).

The larger box size used in the reconstructions allowed for a larger field of view to display all or partial densities for ∼77 flagellin subunits per map, covering 1-3/11 helical turns along the 5-start axis for FljK (378.7 Å) and FljL (377.2 Å), and 1-4/11 helical turns for FljM (381.5 Å) (Fig 1). The flagellin packing for the three reconstructions had a right hand 11-start angle (Fig. 2). However, the FljM D1 domain appeared parallel to the helical axis (Fig. 2A), while the D1 domain in FljK and FljL appeared slightly rotated to the left, when compared to the helical axis. A cross-section slice of the volume showed a clear central lumen of 25.6 Å, 25.5 Å, and 24.3 Å for FljK, FljL, and FljM filaments, respectively (Fig. 1). The filament diameter was similar for all three maps and displayed an outer diameter of 133.5 Å, 131.9 Å, and 134.4 Å for FljK, FljL, and FljM filaments, respectively (Fig. 1).

A slice along the helical axis revealed the highest resolved regions were along the central part of the maps (Fig. 1C, F, and I), because helical reconstruction methods focus particle alignment along a defined percentage of the helical axis. The FljM map was best resolved, with well-defined sides chains even at the far ends of the filament map, most likely due to the rigidity of the FljM filament structure. Finally, the cross-section perpendicular to the helical axis revealed that surface exposed regions were slightly less resolved in all 3 maps when compared to the central regions of the flagellins, but these regions resolved to below 3.5 Å in each model. The symmetrized maps were used for model building.

### Asymmetrical Helical Reconstruction and 3D Variability Analysis

Historically, helical reconstruction methods were applied to straightened filaments and structural analysis was restricted to these locked confirmations. However, new processing tools alleviate the need for straightening mutations to study helical polymers. Qualitative analysis of filament micrographs show that FljM filaments are straighter than FljK or FljL filaments. To validate this observation a round of asymmetrical helical reconstruction was performed in cryoSPARC. Particles were exported from RELION-4.0 into cryoSPARC followed by a helix refinement job with no helical parameters, and the final RELION map was low pass filtered to 20 Å and used as the initial reference for the asymmetrical reconstruction. The resulting FljL and FljK maps showed curvature along the helical axis while FljM remains rigid (Fig. S3, S5, S7). This FljM map resolved to ∼2.49 Å and was used for building a non-symmetrized model (Table S1). To further improve map quality for the curved FljK and FljL reconstructions, a round of 3D classification was performed followed by asymmetrical reconstruction. These maps resolved to ∼2.84 Å and ∼2.92 Å for FljK and FljL, respectively, and were used for building non-symmetrized models (Table S1). To further analyze filament flexibility a 3D variability analysis (3DVA) was performed. Each particle set was used to solve three clusters or possible states present in the data set. These clusters were then assembled into a movie to visualize subtle differences. As expected, the FljM 3DVA was more rigid with little movement along the length of the flagellum reconstruction, while both FljK and FljL displayed higher levels of flexibility. These subtle differences impacted map quality and resolution, though they were not detrimental to the reconstruction workflow.

### Conserved Glycosylation Sites Reveal Surface Ridges

Filament models were displayed as surfaces and overlayed on the corresponding electron potential map to reveal additional densities in the D1 region (Fig. 3A, 3D, 3G). These additional densities were assumed to be glycosylated threonine residues (Fig. 3C, 3F, 3I). Analysis of individual flagellin subunits revealed these glycosylation sites were conserved across the three flagellin types and were uniquely arranged to form ridges about 40 Å apart, spanning the entirety of the reconstruction (Fig. 3). These glycosylation sites were located near the β-hairpin region, comprised of residues 145 to 160 in FljK and FljM, and residues 144 to 159 in FljL, which was critical for flagellin packing and switching. Threonine 143 and 163 were located just before and after the β-hairpin, respectively, while threonine 158 was located on the c-terminal β-strand of the β-hairpin (Fig. 3B, 3E). Threonine 196 was located on the D1 c-terminal α-helix which was adjacent to the β-hairpin region (Fig. 3B, 3E). Finally, threonine 107 appeared to be located distally to this region, but upon further inspection this residue was located between threonine 143, 158, and 163 of the adjacent *N+11* and *N+5* flagellins (Fig. 3B, 3E). The locations of these sites were conserved in FljL with each position shifted by one residue, e.g., threonine 107, 143, 158, 163, and 196 in FljM and FljK corresponded to threonine 106, 142, 157, 162, and 195 in FljL, respectively (Fig. 3H).

**Figure 3.**
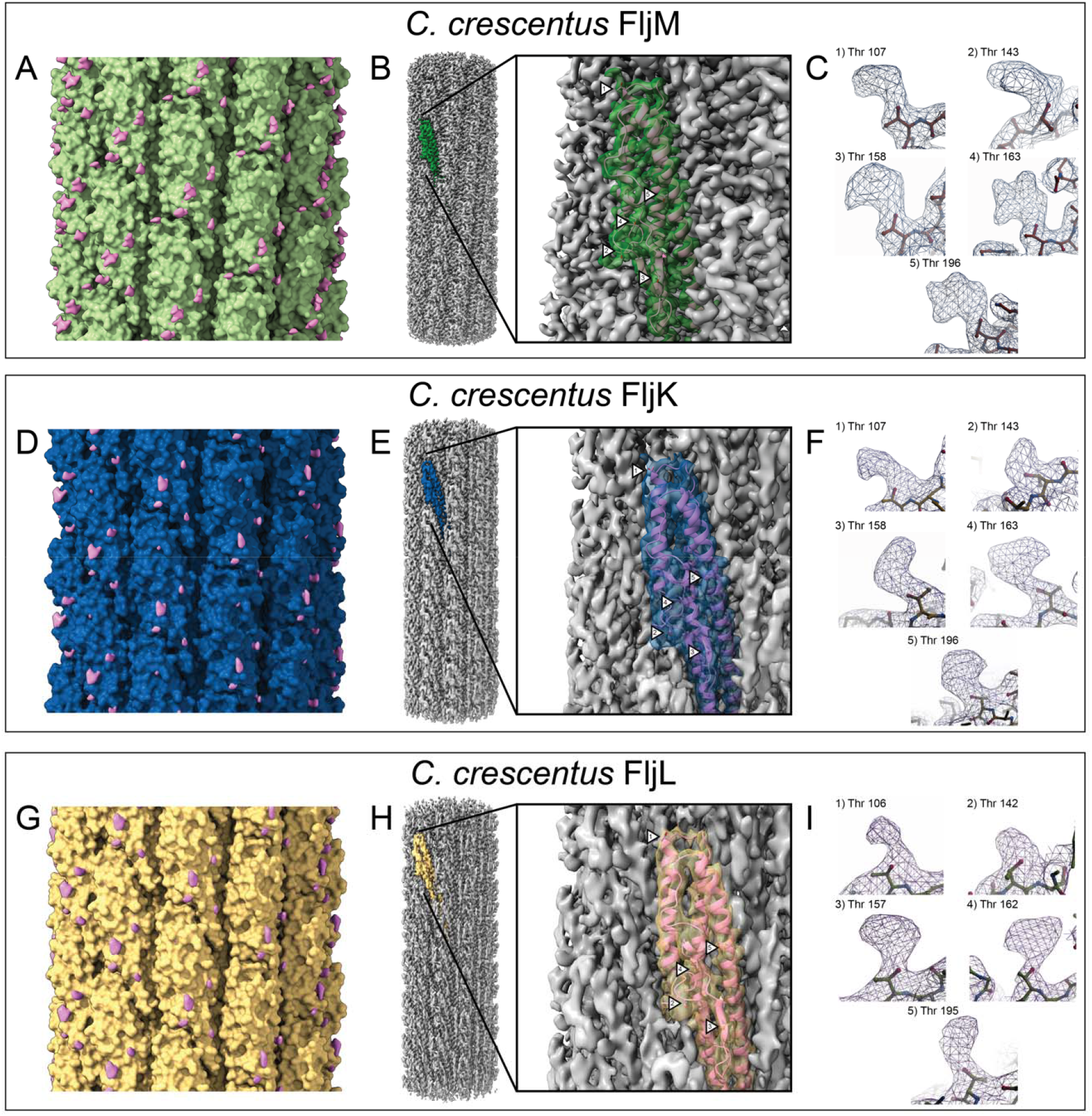
Glycosylation sites on the flagellin D1 domain form ridges along the 6-start angle. (A, D, G) Surface representations of the FljM (A, green), FljK (D, blue), and FljL (G, yellow) filament models overlayed on the electron potential map (pink) reveal extra densities (glycosylations) on the D1 domain. (B, E, H) Segmentation of a single FljM (B, green), FljK (E, blue), and FljL (H, yellow) flagellin subunit along the helical axis and closeup of the electron potential map with the corresponding atomic model (pink). Arrows indicate conserved glycosylated threonine residues. (C, F, I) Electron potential maps (gray transparent) with extra density at threonine residues. Numbering corresponds to arrows in B, E, and H.

### A Network of Hydrogen Bonds Along the *N, N+5, and N+11* Interface Stabilize Flagellin Packing and Provide Evidence for Multi-flagellin Filament Stabilization

The wild-type *C. crescentus* filament is comprised of up to six flagellin subunits, thus conserved residues are key to stabilizing flagellin-flagellin interactions regardless of the subunits involved. Analysis of the D0 domains between a flagellin, *N,* (Fig. 4A, 4E, 4I, green) and the adjacent subunit along the 5-start, *N+5,* (Fig. 4A, 4E, 4I, tan) and 11-start subunit, *N+11,* (Fig. 4A, 4E, 4I, pink) revealed a network of hydrogen bonds between the three subunits at this interface. FljM had the most compact network of hydrogen bonds, of the three *Caulobacter* flagellins examined here, due to a smaller translational displacement along the 11-start angle (Fig. 2A). Asparagine 7 of the *N+11* flagellin acted as an anchor to interact with lysine 250 of the *N+5* subunit, asparagine 234 of the *N* flagellin, and the backbone carbonyl of threonine 35 of the *N* flagellin (Fig. 4B). There were additional bonds between *N+5* lysine 250 and *N* threonine 35, and *N* asparagine 234 and the backbone nitrogen of *N+11* threonine 8 (Fig. 4B). The increase in translational displacement along the 11-start angle for FljK and FljL, when compared to FljM (Fig. 2), resulted in a loss of hydrogen bonds to the *N* flagellin backbone at this interface. However, the changes in rotational displacement along the 11-start angle resulted in additional hydrogen bonds with the N-terminal and C-terminal α-helices of the *N+5* subunit. In FljK, the *N+11* asparagine 7 again acted as an anchor to interact with *N+5* lysine 250 and *N* aspartate 234 (Fig. 4F). However, new side-chain interactions between *N* threonine 35 and *N+5* asparagine 18, *N* lysine 37 to *N* asparagine 234, and *N+5* serine 246 to the *N+11* isoleucine backbone nitrogen were noted (Fig. 4F). In FljL, which had the lowest motility of the three mono-flagellin filaments, there was a network of only three hydrogen bonds at this site. Again, *N+11* asparagine 6 interacted with *N+5* lysine 249 and *N* aspartate 233, and *N* asparagine 233 interacted with *N* lysine 36 (Fig. 4J). There were no additional hydrogen bonds to the FljL *N+5* flagellin, as observed in the FljK filament. Overall, the slight changes in helical twist and rise resulted in a reorganization of the *N, N+5, N+11* interface, which acted as a hinge that changed the global architecture of the filament.

**Figure 4.**
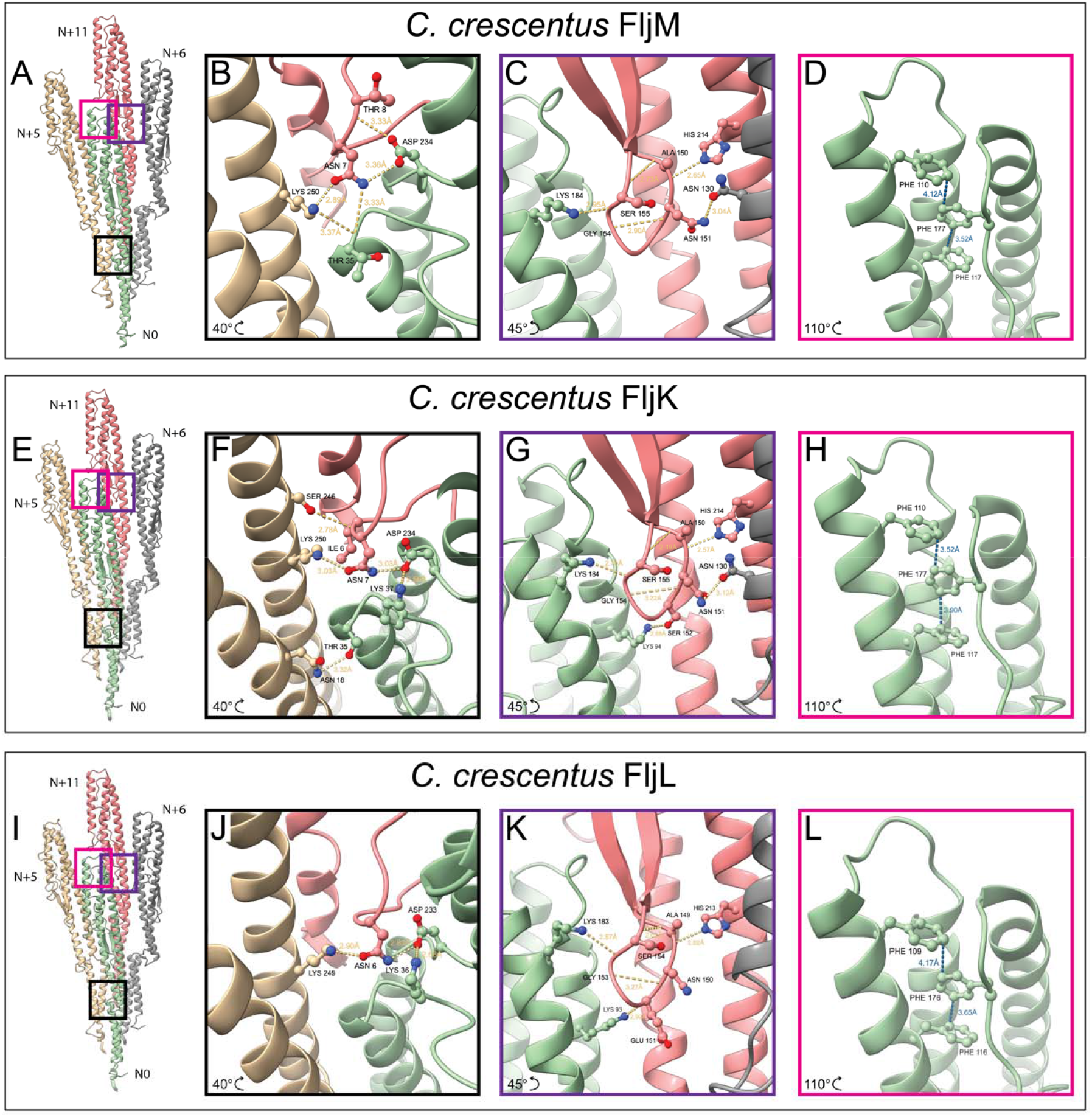
*C. crescentus* flagellin interactions stabilize the flagellar filament and conserved residues are key for multi-flagellin filament formation. (A, E, I) Flagellin subunit (*N0*, green) is stabilized by interactions with neighboring subunits (*N+5*, tan, *N+*6, gray, *N+11*, pink). (B, F, J) The hinge region in the FljM (B), FljK (F), and FljL (J) filament is stabilized by a network of hydrogen bonds (yellow dashed lines) between subunits *N+5* (tan) and *N+11* (pink). (C, G, K) For filaments FljM (C), FljK (G), and FljL (K), the β-hairpin of subunit *N+11* (pink) is stabilized by intra-flagellin interactions and a network of hydrogen bonds with subunits *N0* (green) and *N+6* (gray). (D, H, L) For all three filaments, pi-stacking (blue dashed lines) interactions between conserved phenylalanine residue stabilize the flagellin tertiary structure.

### The Flagellin **β**-Hairpin is Critical for Filament Quaternary Structure

Similarly, to the *N, N+5, N+11* interface, the β-hairpin region was restructured due to subtle rotational and translational shifts between flagellins. In all three models, the loop of the *N+11* subunit was stabilized by backbone interactions between alanine to serine, and asparagine to glycine (Fig. 4C, 4G, 4K). There was a key intra-flagellin histidine located on the D1 C-terminal α-helix that formed a hydrogen bond with the carbonyl group of the loop alanine, residue 150 in FljK and FljM, and residue 149 in FljL (Fig. 4C, 4G, 4K). Additionally, all three filament models contained a hydrogen bond between the *N0* lysine, which is lysine 184 for FljK and FljM, and lysine 183 for FljL, to the *N+11* loop glycine carbonyl, residue 154 in FljK and FljM, and residue 153 in FljL (Fig. 4C, 4G, 4K). These conserved interactions provided evidence for how the key contacts likely supporting multi-flagellin interactions that form stable polymers without the need for additional accessory proteins or stabilizing platforms between different flagellin types. FljM and FljK formed additional side chain interactions between *N+11* asparagine 151 and *N+6* asparagine 130 (Fig. 4C, 4G, gray chain), however this interaction was not present in FljL. Finally, FljK formed an additional hydrogen bond between *N0* lysine 94 and *N+11* serine 152 of the β-hairpin loop, while a similar bond occurred in FljL between the *N0* lysine 93 and the *N+11* backbone carbonyl of glutamine 151 (Fig. 4G, 4K). In the hinge region, the slightly smaller translational displacement between FljM subunits resulted in a compact region with additional bonds between the 11-start subunits, as compared to the FljK and FljL models. However, in the β-hairpin region the smaller rotational displacement observed in FljK resulted in an additional hydrogen bond between the loop and the adjacent flagellins, when compared to FljL and FljM. Overall, all three models contain conserved residues that stabilized these regions and could support multi-flagellin filament formation.

### Pi-Stacking Interactions in the D1 Domain Support Tertiary Flagellin Structure

*C. crescentus* flagellins do not contain D2 or D3 domains as observed in other bacterial systems (26, 30, 31). Instead, the D1 domain serves as the surface exposed region of the *Caulobacter* flagellin monomer. In the three models presented here, the tertiary structure of the D1 domain was supported by a set of three phenylalanine residues with pi-stacking interactions (Fig. 4D, 4H, 4L). Specifically, in FljK and FljM, phenylalanine 177 was sandwiched in between phenylalanine 110 and 117, which were both perpendicular to the central ring forming pi-teeing interactions (Fig. 4D, 4H, 4L). These interactions were also seen in FljL between phenylalanine 109, 116, and 176 (Fig. 4D, 4H, 4L). The two former rings were located on the N-terminal portion of the D1 domain, while the latter was located on the C-terminal portion of the D1 domain and provided a support for the flagellin monomer.

### Filament Synthesis and Functionality Examined via Negative Stain Analysis and Motility Assays

Whole cell negative stain imaging was performed to validate the presence of intact flagellar filaments in the *C. crescentus* swarmer cells. All three mono-flagellin strains synthesized filaments with varying lengths. Strains will be referred to by the flagellin gene present in the strain. FljL displayed the shortest filament measuring 0.9 μm, the FljK filament was 3.4 μm in length, and FljM filaments were 4.2 μm in length (data not shown here). Wild-type filaments averaged 5.9 μm in length, while the no flagellin strain did not produce a filament. Filament functionality was assessed by motility assays. Bacterial growth was monitored over 72 hours and values were displayed as a percentage of wild-type growth, where wild-type growth was 100% (Fig. S1). All single flagellin filaments assessed here showed severely reduced levels of motility. The FljK strain had the highest motility measurement at ∼52%, followed by FljM at ∼32%, FljL at ∼27%, and the no flagellins strain at ∼16%. The reduction in motility observed in FljL appeared to be tightly related to filament length, however FljM possessed a much longer filament and slightly higher motility. This difference seemed to be related to the subunit packing in the FljM filament leading to a straighter filament as compared to FljK and FljL.

## Discussion

The latest advancements in predicting protein tertiary structure using AlphaFold2 (32) has accelerated protein structure determination. However, these predications lack essential quaternary structural information. Here we used these predications as a basis for structure determination by refining a predicted model into our cryo-EM electron potential density maps, followed by docking and refinement of additional chains throughout the map (Fig. 1, 2). All three symmetrized cryo-EM structures determined here resolved to below 2.7 Å, and all three asymmetrical reconstructions resolved to below 3.0 Å. All electron potential maps allowed us to build atomic models with clear side chain densities for 44 subunits per map. The resolution and size of these reconstructions provided clear evidence surrounding the molecular interactions that impact flagellin packing and how they influence the filament ultrastructure. Additionally, our models provided clues as to how the multi-flagellin wild-type filament may form a continuous structure without accessory proteins to support variation between the flagellin monomers.

We note that the hinge region and β-hairpin form a network of hydrogen bonds that may regulate flagellin packing and the overall architecture of the bacterial flagellum. Conserved residues at both sites provide essential contacts for neighboring flagellin subunits that support their quaternary structure. It seems that histidine 214 in FljK and FljM, or histidine 213 in FljL, is involved in stabilizing the quaternary structure of a filament comprised of multiple flagellins. Similarly, asparagine 7 in FljK and FljM, or asparagine 6 in FljL, may act as an anchor in the hinge region of the wild-type filament providing support regardless of the identity of the neighboring flagellin subunit. We observed that, at the tertiary structure level, the three flagellins are homologous, but small changes at the amino acid level result in minor rotational and translational shifts between subunits that impacts the overall architecture of the filament.

The quaternary structure revealed that repeating glycosylation sites, across all three filaments, results in a ridge along the 6-start subunits (Fig. 3). These glycosylation sites are just before and after the β-hairpin and may provide protection for this surface exposed region. Removal of these glycosylations or mutations to these threonine residues may elucidate whether they are critical for motility, filament stability, or other functionalities, and will be a key goal of future work.

Understanding the role of these conserved molecular contacts in their energetically favorable states provides a structure-guided method for constructing new peptide polymers with unique architectures and different mechanical properties. Developments with peptide-based polymers have focused on the *de novo* design of either α-helical or β-sheet assemblies, with tunable oligomeric states that govern a range of functional properties (33–35). Many of these peptide-based polymers assemble to form nanotubes that are designed to mimic functions of materials found in nature, such as adherence and sensing, delivery, or motility. In the case of an engineered material designed to imitate the bacterial flagellum, the central lumen of an assembly could serve as a reservoir for drug delivery or other biotech applications. Additionally, the amino acid sequence could be modified to synthesize dimorphic filaments that form a solid glass-like state without the need for chemical cross-linkers (33). Alternatively, the sequence governing filament assembly and function could be optimized to support propulsion of synthetic nanostructures (34, 36). Here, our results provide new structural insights on the assembly of helical polymers, offering a basis for rationally engineered soft materials.

## Materials and Methods

### Bacterial Strains, Plasmids, and Growth Conditions

The bacterial strains and plasmids for this study are summarized in Tables S2 and S3. PCR primers for this study are summarized in Table S4. *E. coli* strain DH5[1] was used for plasmid maintenance. The gene for FljM including ∼500 bp upstream, to conserve the native promoter, was amplified and inserted into the pMR10 plasmid, which is kanamycin (50 µg/mL) resistant. Individual isolates were verified by analytical DNA digest and by whole plasmid sequencing for validation (Plasmidsaurus, Eugene, OR). The FljM plasmid was transformed into electrocompetent *C. crescentus* Δ*fljJKLMNO* (no flagellins) and stored in 10% DMSO. Unless otherwise noted, *E. coli* strains were grown in LB medium (BD, Franklin Lakes, NJ) at 37°C and *C. crescentus* strains were grown in peptone yeast extract (PYE; 0.2% peptone, 0.1% yeast extract) supplemented with 1mM MgSO_4_ and 0.5 mM CaCl_2_ at 30°C. Plating medium contained 1.5% (w/v) Bacto agar.

### Motility Assays

Motility assays were conducted on PYE soft-agar plates (motility plates), containing 0.3% (w/v) agar, to test for motility phenotypes as previously described (13). Briefly, three isolated colonies were selected for each strain and grown to an OD_600_ of 0.3. Motility plates with appropriate antibiotics were inoculated with a 1 µL drop of 0.3 OD_600_ culture. Each plate contained four inoculation sites and three plates were prepared for each bacterial strain, one for each isolated colony. Motility plates were incubated at 30°C and imaged every 24 hours for 72 hours. Motility plates were imaged on and iBright CL1500 system (Thermo Fisher Scientific, Hillsboro, Oregon) and evaluated with ImageJ (37). Values are presented as a percentage of motility of the wild-type strain NA1000.

### Flagellar Filament Isolation

*Caulobacter crescentus* flagellar filaments were prepared as previously published (13). Briefly, *C. crescentus* cells were grown to an OD_600_ of 0.6. Cells were pelleted at 10,000 x *g* for 15 minutes. The supernatant was collected and centrifuged at 48,000 x *g* for 35 minutes to pellet flagellar filaments. The pellet was covered with 10 mL of phosphate buffered saline (PBS) and incubated overnight at 4°C with shaking at 100 rpm. Each centrifugation step was repeated to remove additional contaminants followed by overnight incubation in 1 mL of PBS with gentle shaking. The flagella suspension was centrifugated at 17,500 x *g* for 15 minutes. The supernatant was collected, and the centrifugation step was repeated followed by centrifugation of the supernatant at 21,000 x *g* for two hours. The final filament pellet was resuspended in 50 μL of PBS followed by overnight, static incubation at 4°C. Sample quality was analyzed by negative stain EM.

### Grid Preparation

Cryo-EM samples were prepared by depositing four (4) microliter (μL) aliquots of the purified flagella onto glow discharged, R2/1 200 mesh, copper Quantifoil grids (Quantifoil Micro Tools GmbH, Germany) prior to were plunge freezing in liquid ethane using a Vitrobot Mark IV (Thermo Fisher Scientific, Hillsboro, Oregon). Blot times ranged from 2 to 4 seconds, blot forces ranged from 0 to 6 with 0.5 seconds of drain time, and 60 seconds of wait time.

### Cryo-EM Data Collection

Imaging was performed on a Titan Krios G3i FEG-TEM (Thermo Fisher Scientific, Hillsboro, Oregon) operated at 300 kV, and equipped with a Gatan K3 direct electron detector and BioQuantum energy filter (Gatan, Inc., Pleasanton, CA). Dose-fractionated micrographs were collected in correlated-double sampling (CDS), counting mode over a defocus range of -0.5 to -2.5 μm with increment steps of 0.25 μm at a pixel size of 0.834 Å with a total dose of 45 e^-^/Å^2^ (1 e^-^/Å^2^/frame). Movies were acquired with 3 shots per hole acquiring multiple holes per stage position by using EPU/AFIS (Thermo Fisher Scientific, Hillsboro, Oregon), on average ∼250 movies were collected per hour.

### Helical Reconstruction

Helical reconstruction was conducted within the RELION-4.0 framework (29). Gain-normalized micrographs were motion corrected with MotionCor2 (38) and initial estimates for whole-micrograph defocus values was performed with gCTF (39). A random subset of 25 micrographs were used for manual picking of filament segments. Particles were extracted with a box size of 500 pixels, rescaled to 250 pixels, and an inter-box distance of 50 Å with an initial helical estimate of 4.8 Å. 2-D classification was performed using a T-regularization value of 2, 25 classes, 200 VDAM mini-batches, and a mask diameter of 350 Å. The best 2D classes were selected and the particles from these classes were used as templates for Topaz (40) training using the Topaz wrapper within the RELION-4.0 Auto-picking job. Additionally, the standard topaz executable was replaced with the Topaz-filament executable which supports start-end coordinate picking for subsequence helical reconstruction in RELION. After 10 iterations of training, the resulting model file was used to perform Topaz picking on a 2^nd^ subset of 20 micrographs with a particle diameter of 400 Å, the additional Topaz argument for filament picking (-f) was supplied and several picking thresholds (-t) ranging from 0 to -6 were tested. A picking threshold of -2 resulted in an average of 10 segments per micrograph and manual inspection of the micrographs confirmed picking along flagellar filaments. These parameters were supplied for automated picking on our entire data set of 8,813 micrographs for FljK, 3,780 micrographs for FljL, and 9,780 micrographs for FljM. Particles were extracted to a particle box size of 500 pixels and re-scaled to 250 pixels followed by 2-D classification using a T-regularization value of 2, 100 classes, 200 VDAM mini-batches, and a mask diameter of 350 Å. Particles contributing to the best classes were then re-extracted and re-centered.

Initial estimates for helical symmetry were taken from homologous flagellins of known structures (13). 3D classification was performed with an initial helical rise of 4.8 Å and helical twist of 65.6°, with local searches of symmetry spanning ± 0.5 Å for helical rise and ± 1° for helical twist. A featureless cylinder was generated using *relion_helix_toolbox* and was used as an initial reference for 3D classification with a T-regularization value of 4, 20 classes, and an angular sampling interval of 3.7°. The resulting volumes were manually inspected and classes showing clear secondary structure features that are consistent with previously determined flagellin structures were selected for an additional round of 2D and 3D classification. After each round of 3D classification, 11-start angles were calculated and classes with clear secondary structures converged to similar 11-start angles ± 0.5°. After the final round of 3D classification one volume was lowpass filtered to 20 Å and used as an initial reference for helical auto-refinement with optimization of helical symmetry parameters for the best classes. The 3D auto-refinement models reached the theoretical limit for 2x binned data at which point particles were re-extracted to the native pixel size (0.834 Å) followed by per-particle CTF refinement and Bayesian polishing before a final round of 3D auto-refinement, followed by masking and post-processing. The final polished particle set was then imported into cryoSPARC for an additional round of 3D refinement with applied helical symmetry.

### Asymmetrical Reconstruction

RELION Bayesian polished, or shiny, particles were imported into cryoSPARC (28) for further processing and analysis. First, we ran a helical refinement job without imposing helical symmetry and using the final RELION map, low pass filtered to 20 Å, as an initial reference. Next, the Symmetry Search Utility was applied to our asymmetrical map to validate helical parameters, before performing a helical refinement job with imposed symmetry followed by local resolution estimates. For the FljK and FljL models, after generating an initial asymmetrical map, a round of 3D classification was performed followed by asymmetrical refinement, symmetry search, helical refinement with imposed symmetry and local resolution estimates.

The particles and resulting mask from the initial asymmetrical helical refinement job were down-sampled 2-fold to a pixel size of 1.668 Å and used as inputs for a 3D variability analysis. The resulting 3D variability components and particles were used as inputs for a 3D variability display job with the following specialized parameters: cluster output mode, 3 clusters, and a filter resolution of 5 Å.

### Model Building and Validation

To accelerate model building, we supplied the corresponding amino acid sequences to Alphafold2 (32) to generate a structure predication model of the appropriate flagellin monomer. The predicted model was fitted into the cryo-EM map using ChimeraX (41, 42), followed by real space refinement with Ramachandran (43) and secondary structure restraints in Phenix (44). The refined monomer was manually placed throughout the cryo-EM map followed by refinement in Phenix until convergence. The final models exhibit excellent stereochemistry as reported in Phenix. Additionally, Phenix reports map to model correlation coefficients of 0.78, 0.78, and 0.88 for FljK, FljL, and FljM, respectively, serving as a cross-validation metric for map quality and helical symmetry parameters.

## Acknowledgments

This work was supported in part by the University of Wisconsin, Madison, the Department of Biochemistry at the University of Wisconsin, Madison, and public health service grants R01 GM104540 and U24 GM139168 to E.R.W. from the NIH. This work was supported in part by the U.S. Department of Energy, Office of Science, Office of Biological and Environmental Research under Award Numbers DE-SC0018409. J.C.S. was supported in part by the Biotechnology Training Program at the University of Wisconsin, Madison, T32 GM135066. D.P. was supported in part by F32 GM143854 from the NIH. We are grateful for the use of facilities and instrumentation at the Cryo-EM Research Center in the Department of Biochemistry at the University of Wisconsin, Madison. We are grateful for the computational resources supplied through the SBGrid Consortium (45)

## Author Contributions

E.R.W. and J.C.S. designed research; J.C.S., E.J.M., N.T.P., D.P., J.Y., B.S., and K.C. performed research; J.C.S. contributed new reagents/analytic tools; J.C.S., J.K.B., and E.R.W. analyzed data; and J.C.S. and E.R.W. wrote the paper.

## Data deposition

The atomic coordinates have been deposited at the Protein Data Bank under accession numbers XXX, XXY, YYY, YYZ, ZZZ, and ZZA. The density maps have been deposited at the Electron Microscopy Data Bank under accession numbers EMD-XXX, EMD-XXY, EMD-YYY, EMD-YYZ, EMD-ZZZ, and EMD-ZZA.

## Supplemental Materials

**Table S1.** Data collection, asymmetrical reconstruction in CryoSPARC, and model statistics.

| Protein | CcFljK | CcFljL | CcFljM |
| --- | --- | --- | --- |
| PDB ID | XXY | YYZ | ZZA |
| EMDB ID | EMD-XXY | EMD-YYZ | EMD-ZZA |
| **Data collection and processing** |  |  |  |
| Voltage (kV) | 300 | 300 | 300 |
| Electron exposure (e<sup>-</sup>/Å<sup>2</sup>) | 45 | 45 | 45 |
| Number of frames | 45 | 45 | 45 |
| Nominal defocus range (-μm) | 0.5 - 2.5 | 0.5 - 2.5 | 0.5 - 2.5 |
| Pixel size (Å) | 0.834 | 0.834 | 0.834 |
| Symmetry (beyond helical) | C<sub>1</sub> | C<sub>1</sub> | C<sub>1</sub> |
| Micrographs | 8,813 | 3,780 | 9,780 |
| Initial particles | 3,672,603 | 2,061,739 | 1,007,292 |
| Final particles | 106,162 | 104,598 | 248,115 |
| Box size (Å) | 417 | 417 | 417 |
| Box size (pixels) | 500 | 500 | 500 |
| Helical symmetry |  |  |  |
| rise (Å) | NA | NA | NA |
| twist (degrees) | NA | NA | NA |
| 11-start filament hand | NA | NA | NA |
| Helical Z parameter (%) |  |  |  |
| masking | 85 | 85 | 85 |
| refinement | 30 | 30 | 30 |
| Map resolution (Å) | 2.84 | 2.92 | 2.49 |
| FSC threshold | 0.143 | 0.143 | 0.143 |
| Sharpening B factor (Å<sup>2</sup>) | -36 | -41 | -63 |
| **Structure Refinement** |  |  |  |
| Total atoms | 85,404 | 86,064 | 85,228 |
| ligands | 0 | 0 | 0 |
| water | 0 | 0 | 0 |
| RMS, bonds (Å) | 0.003 | 0.003 | 0.013 |
| RMS, angles (°) | 0.778 | 0.750 | 0.741 |
| Ramachandran favored (%) | 97.79 | 97.73 | 98.37 |
| Ramachandran outliers (%) | 0.21 | 0.28 | 0.04 |
| Rotamer outliers (%) | 0.01 | 0.05 | 0.01 |
| Map-model correlation (masked) | 0.82 | 0.85 | 0.94 |
| Model B factors |  |  |  |
| minimum | 43.41 | 52.46 | 44.68 |
| mean | 98.68 | 109.74 | 88.89 |
| maximum | 325.37 | 350.89 | 183.94 |
| MolProbity score | 1.69 | 1.73 | 1.18 |
| Clashscore | 13.62 | 14.53 | 3.93 |

**Table S2.** *Caulobacter crescentus* strains used in this study. All strains are derived from NA1000.

| Organism | Strain | Genotype | Description | Reference |
| --- | --- | --- | --- | --- |
| *C. crescentus* | NA1000 | Wild type | *C. crescentus* wild-type strain | (46) |
| *C. crescentus* | TPA2357 | $\Delta fljJ \Delta fljK \Delta fljL \Delta fljM - fljO$ | Strain with a deletion for all flagellin genes | (15) |
| *C. crescentus* | TPA2353 | $\Delta fljJ \Delta fljL \Delta fljM - fljO$ | Strain with deletions for flagellins other than *fljK* | (15) |
| *C. crescentus* | TPA2346 | $\Delta fljJ \Delta fljK \Delta fljM - fljO$ | Strain with deletions for flagellins other than *fljL* | (15) |
| *C. crescentus* | ERW2301 | $\Delta fljJ \Delta fljK \Delta fljL \Delta fljM - fljO$ | TPA2357 carrying pERW2301 encoding *fljM* | This study |

**Table S3.** *Caulobacter crescentus* plasmids used in this study.

| Plasmid | Description | Reference |
| --- | --- | --- |
| pMR10 | Broad-host-range cloning vector, Kan<sup>r</sup>, low copy number | (46) |
| pCS12 | Construct encoding *fljK* T103C N130S | (13) |
| pERW2301 | Construct encoding *fljM* and it's native promoter region | This study |

**Table S4.**
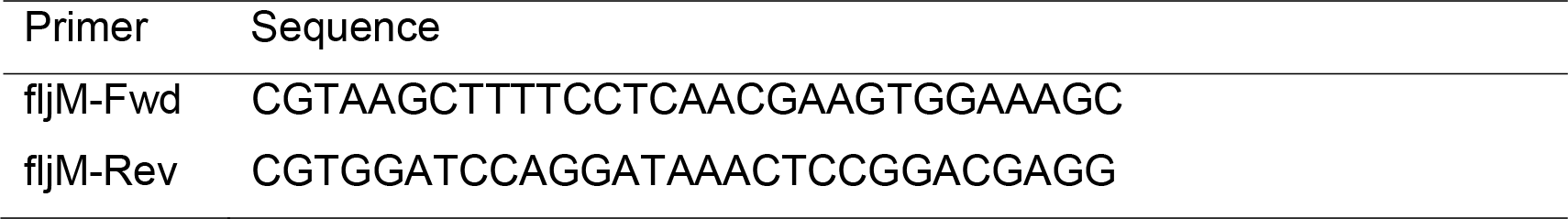
Primers used in this study.

**Figure S1.**
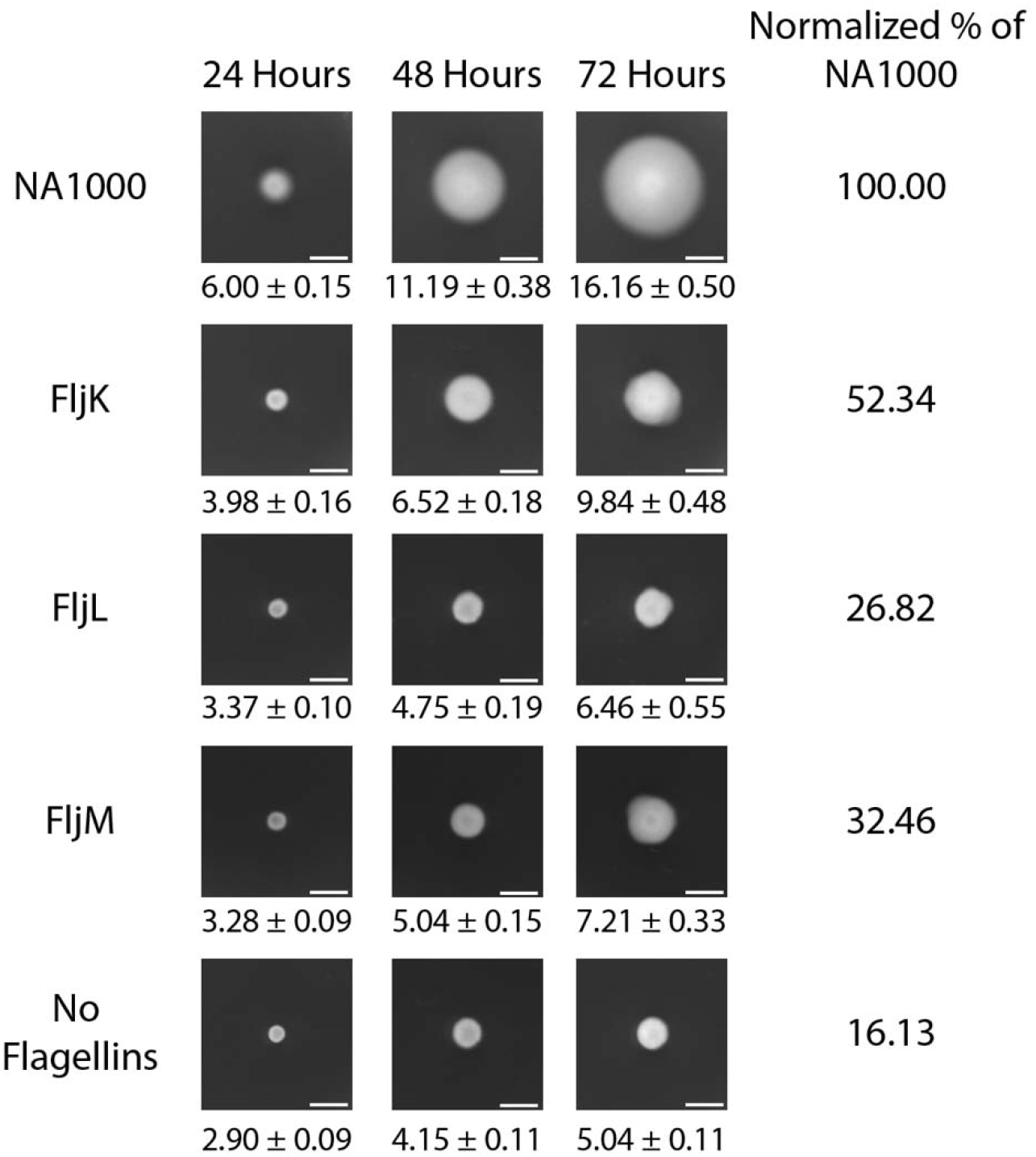
*Caulobacter crescentus* motility assays over 72 hours. Motility of wild-type (NA1000), FljK, FljL, FljM, and no flagellin (ΔFljJKLMNO) strains were assayed on soft agar motility plates. Motility was measured at 24-, 48-, and 72-hours post inoculation. Normalized percentages of motility are displayed as a percentage of wild-type (NA1000) motility. The presented images and normalized percentage of motility values are representative of 3 independent biological replicates. Growth is measured in millimeters (Scale bar, 5 mm).

**Figure S2.**
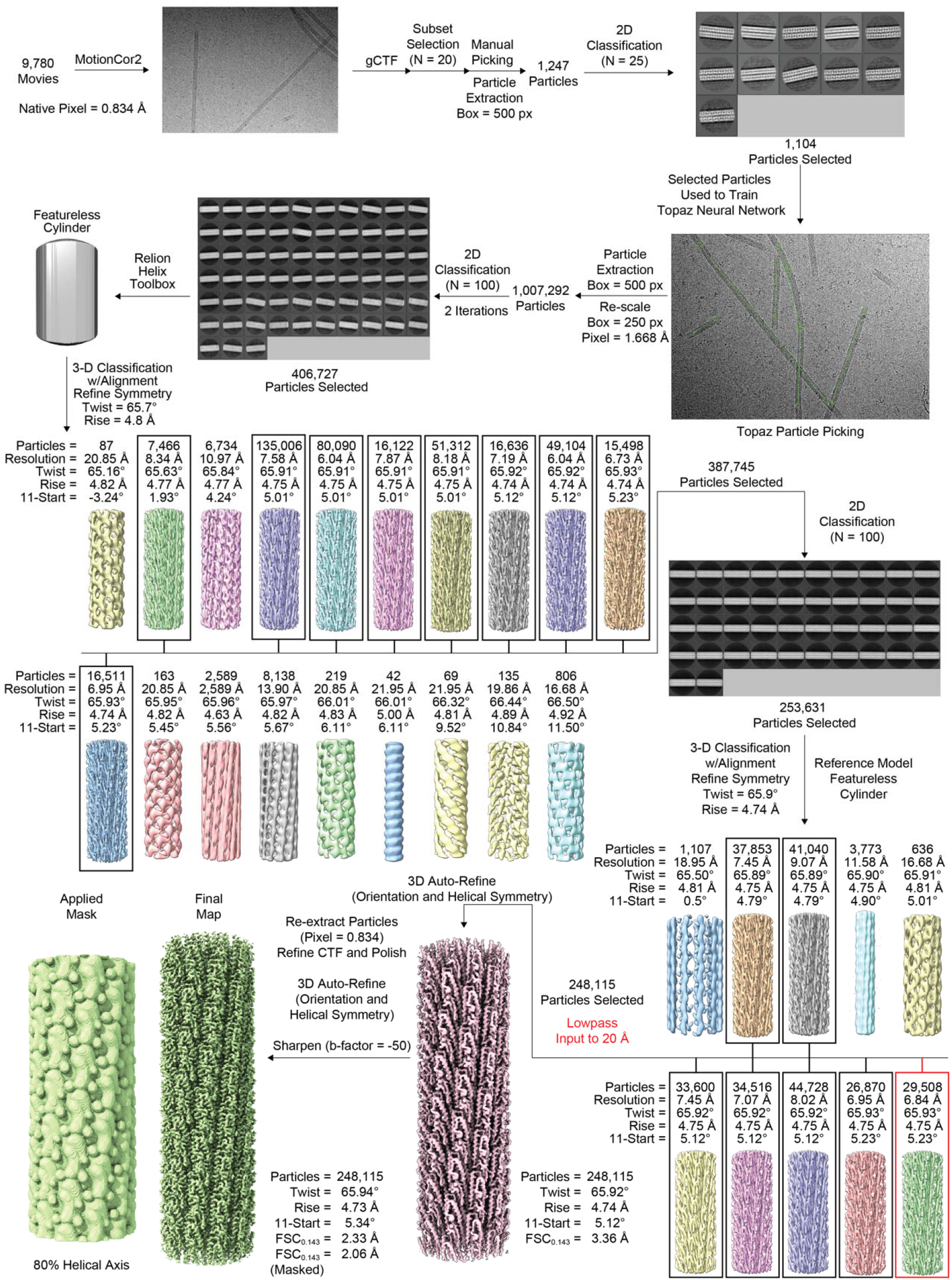
FljM filament RELION workflow. Reconstruction pipeline detailing particle picking, 2D classification, 3D classification, and 3D-refinement steps for the FljM filament electron potential map.

**Figure S3.**
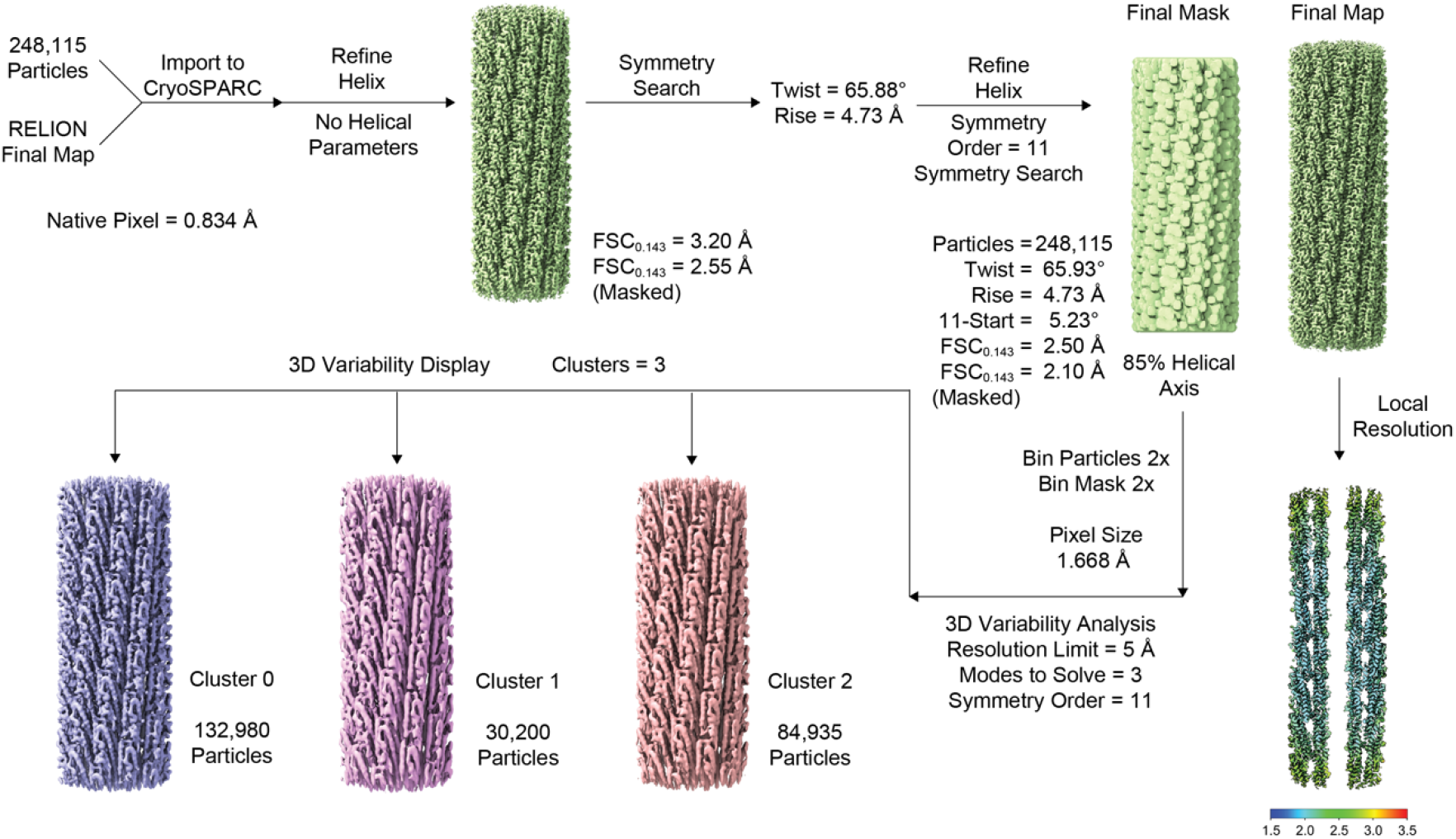
FljM filament CryoSPARC workflow. Additional processing steps in CryoSPARC include asymmetrical helical reconstruction, 3D variability analysis, and local resolution map.

**Figure S4.**
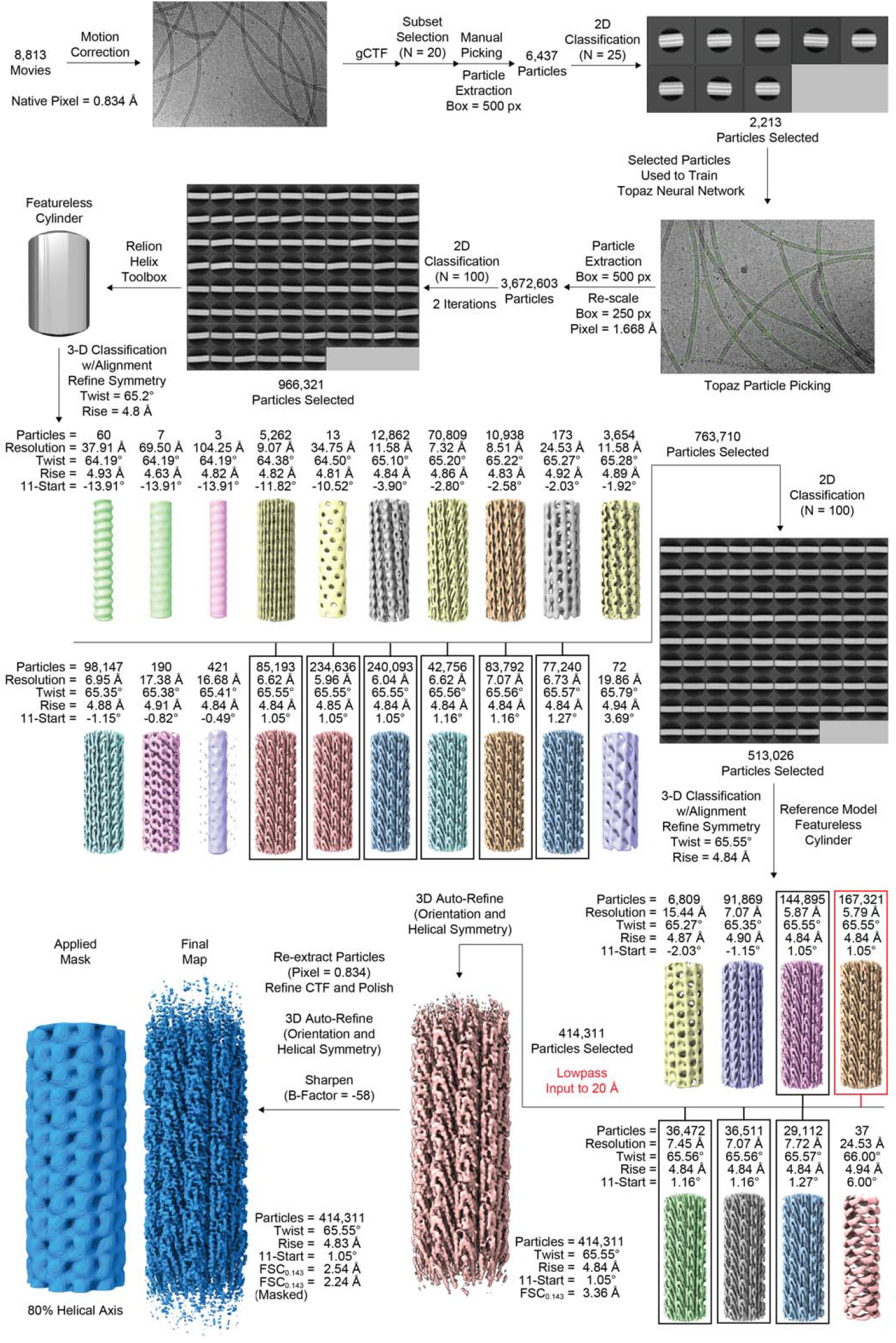
FljK filament RELION workflow. Reconstruction pipeline detailing particle picking, 2D classification, 3D classification, and 3D-refinement steps for the FljK filament electron potential map.

**Figure S5.**
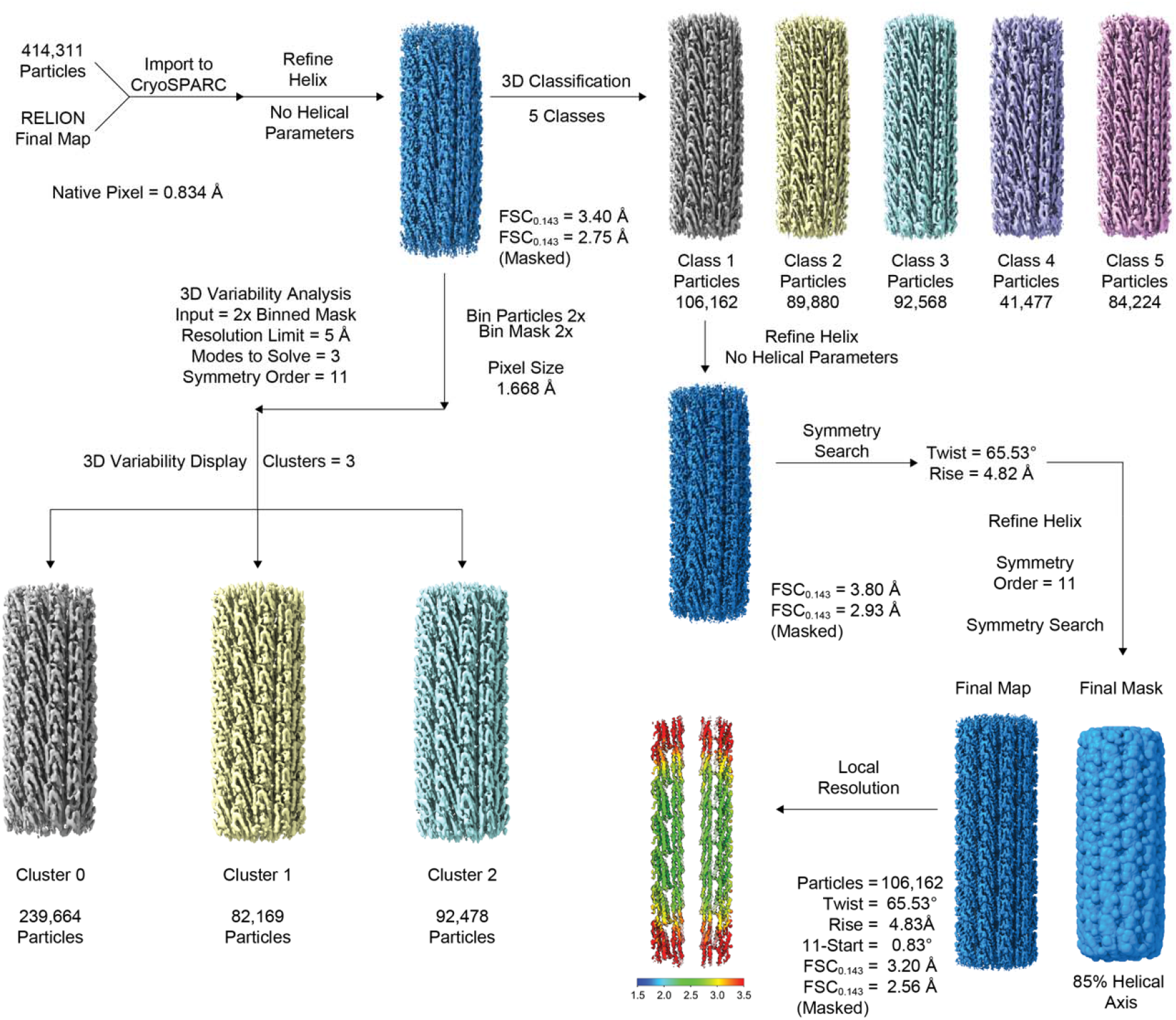
FljK filament CryoSPARC workflow. Additional processing steps in CryoSPARC include asymmetrical helical reconstruction, 3D classification, 3D variability analysis, and local resolution map.

**Figure S6.**
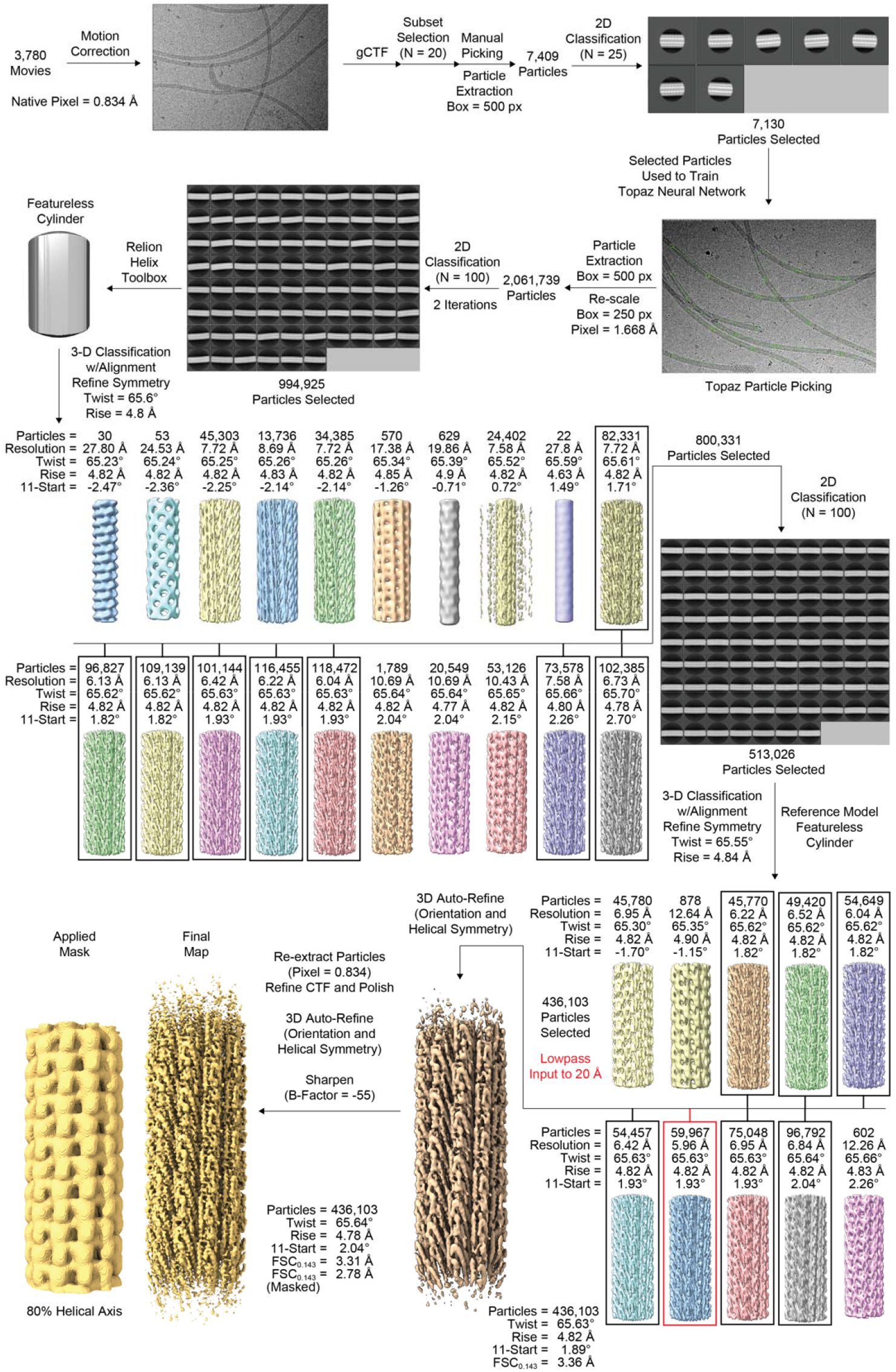
FljL filament RELION workflow. Reconstruction pipeline detailing particle picking, 2D classification, 3D classification, and 3D-refinement steps for the FljL filament electron potential map.

**Figure S7.**
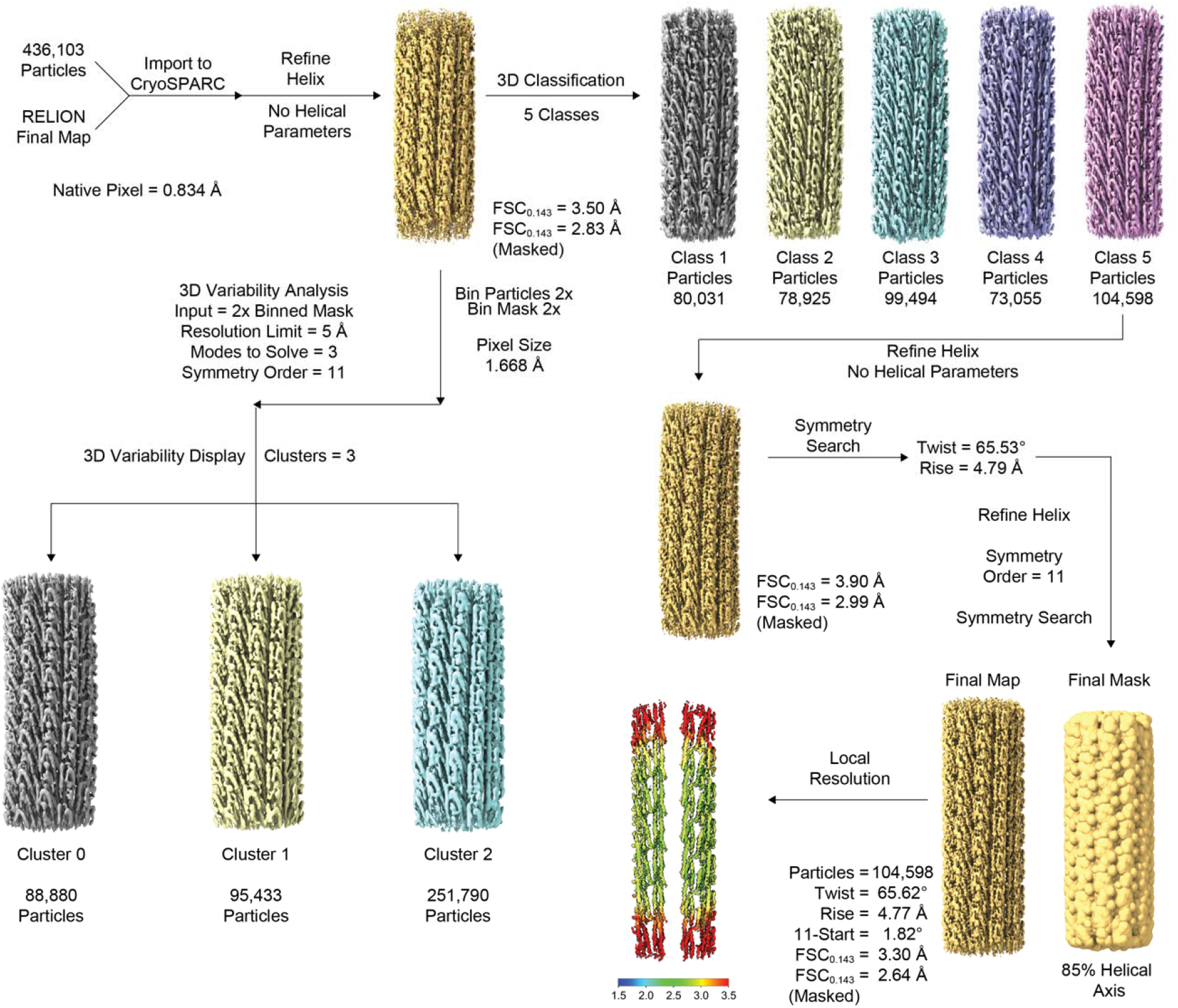
FljL filament CryoSPARC workflow. Additional processing steps in CryoSPARC include asymmetrical helical reconstruction, 3D classification, 3D variability analysis, and local resolution map.

**Figure S8.**
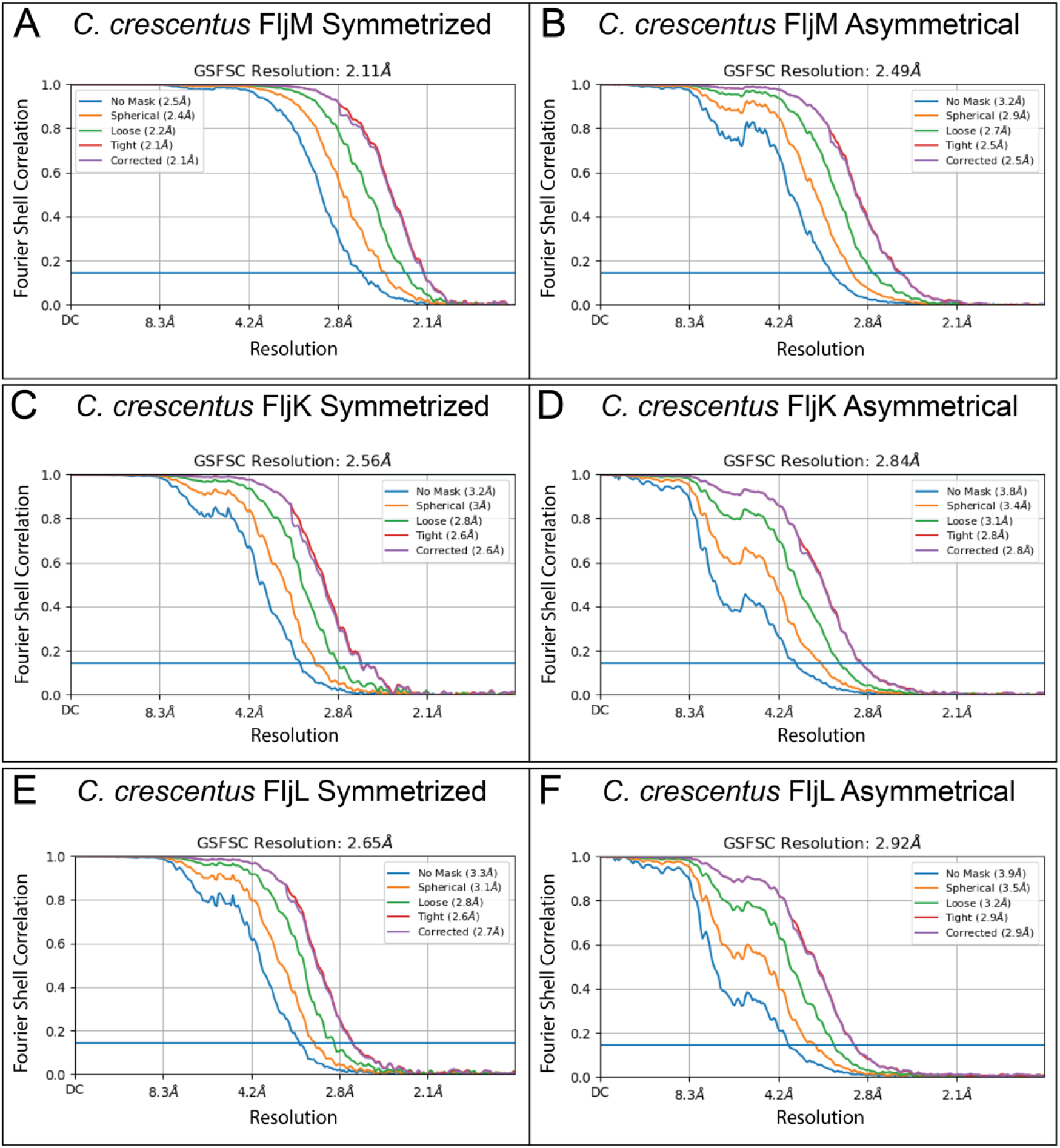
FSC curve comparison. Gold-standard FSC curves generated in CryoSPARC for FljM symmetrized (A), FljM asymmetrical (B), FljK symmetrized (C), FljK asymmetrical (D), FljL symmetrized (E), and FljL asymmetrical (F) reconstructions.

